# New insights into the evolution of Charlie Chaplin worms (Histriobdellidae, Eunicida, Annelida)

**DOI:** 10.1101/2025.10.09.681451

**Authors:** Conrad Helm, Katrine Worsaae, Paul Kalke, Nataliya Budaeva

## Abstract

Histriobdellidae, the so-called Charlie Chaplin worms, is an enigmatic group of microscopic commensal annelids associated with crustaceans. They crawl by alternately attaching their adhesive anterior appendages and left and right huge lateral ‘feet’, and bear a complex jaw apparatus in the ventral muscular pharynx. Although histriobdellids were always thought to be a part of the jaw-bearing clade Eunicida, their exact placement within the annelid tree is still debated due to their highly derived external morphology and long branch attraction artefacts in molecular analyses.

In this study we employ morphological and molecular comparative approaches in order to gain new insights into the evolution of Histriobdellidae and its aberrant traits. Our phylogenetic analyses of Eunicida including 52 species and four molecular markers yield further support for Histriobdellidae being the sister group to the eunicid family Dorvilleidae. The detailed morphology of *Histriobdella homari* Van Beneden, 1858, a commensal of the European lobster, was examined using standard immunohistochemical stainings and subsequent confocal laser scanning microscopy (CLSM) as well as scanning electron microscopy (SEM).

Integrative analyses allow us to compare in detail with other eunicidans and unravel extensive anatomical transformations in Histriobdellidae. Neural innervation patterns help verify the presence of antennae and true annelid palps on the histriobdellid prostomium. The arrangement of ganglia and the neuronal scaffold innervating the anterior end supports the presence of a buccal segment (peristomium) in *Histriobdella*. Additionally, based on our comprehensive investigations we newly propose their adhesive anterior locomotory appendages to be homologous with parapodia, and their posterior-most adhesive locomotory appendages to be homologous with the pygidial lobes of other Annelida.

Detailed studies of this highly deviating family of annelids not only exemplify how to reconstruct extreme transformation of canonical annelid characters such as parapodia, but again also highlight the exceptional evolutionary plasticity of the annelid body plan.

## Introduction

Annelida is a lophotrochozoan taxon, which is well-known for its diversity and morphological variability and their origin dates back to the Cambrian explosion (Eibye-Jacobsen & Vinther, 2012; Purschke, 2016; Parry et al. 2016; Rouse, Pleijel & Tilic, 2022). Annelids occur in a vast range of marine, limnic and terrestrial habitats, and from the deep sea up to the surface of the Arctic glaciers. Consequently, they had to adapt to various environmental conditions and food sources (Purschke, Bleidorn & Struck, 2014). Although always referred to as being segmented and chaetigerous worms (Seaver, 2003), evolutionary transitions and adaptations have led to reductions of segmentation and chaetae in different annelid clades. Hence, enigmatic taxa such as the commensals Myzostomida and Histriobdellidae, the sediment-dwelling Sipuncula as well as the sausage-shaped Echiura (now Thalassematidae sensu Goto et al. (2020)), or the interstitial families Diurodrilidae, Lobatocerebridae, and Dinophilidae represent highly modified adult body plans, which cause difficulties in robust placement of these groups within phylogenetic trees (e.g., Struck et al., 2007; Struck et al., 2011; Weigert & Bleidorn, 2016; Worsaae et al. 2021). Fortunately, the annelid phylogeny became more robust thanks to different phylogenomic approaches in the last years, and many of the so far hardly-placeable aberrant taxa were finally placed within the annelid tree. Nevertheless, several groups are still scarcely investigated and unambiguous phylogenetic placements are still pending (Struck, 2011; Weigert et al., 2014; Andrade et al., 2015; Struck et al., 2015; Laumer et al., 2015; Helm et al., 2018; Martin-Duran et al. 2021; Rouse, Pleijel & Tilic, 2022; Tilic et al. 2022).

The remarkable Charlie Chaplin worms, Histriobdellidae Claus & Moquin-Tandon, 1884, comprise only 13 described species of minute marine and fresh-water commensal annelids that live on decapod and isopod crustaceans (Helm, Vila & Budaeva, 2020). On their respective host, histriobdellid worms mainly populate the branchial filaments as well as the pleopods and egg masses, and can occur in high densities (Lerch & Uglem, 1996). They have direct development and transparent bodies with a minute size of only 0.7–1.5 mm and lacking chaetae. Yet, they can easily be distinguished by a prominent, dark chitinous jaw apparatus and a characteristic posteriormost segment extending laterally into ‘feets’, resembling the laterally pointing, large shoes and walk of Charlie Chaplin (Tzetlin et al., 2020; Rouse, Pleijel & Tilic, 2022). They show resemblance to eunicidan annelids in their jaw apparatus and in carrying five head appendages (Figs. 2a), considered as antennae (three dorsal) and palps (two latero-ventral), respectively (see also Zanol, 2010; Kuhl et al., 2022). Although the general jaw architecture of histriobdellid mandibles and maxillae is similar to that of other eunicidan families (Helm, Vila & Budaeva, 2020), the classification of the maxillary apparatus as one of the known types, i.e. ctenognath (Dorvilleidae), symmetrognath (Lumbrineridae), eulabidognath (Eunicidae and Onuphidae), or prionognath (Oenonidae) is debated (Zanol et al. 2021). Histriobdellid maxillae were described as being typical for Eunicida (Mesnil and Caullery 1922), either of the prionognath (Rouse and Pleijel 2001) or the ctenognath type (Tzetlin 1980, Tzetlin et al. 2020), or unlike any other (Paxton 2009).

The placement of Histriobdellidae within Eunicida has so far been problematic due to their deviating morphology preventing the establishment of primary homologies of the main body parts with those of other eunicidans. Moreover, in molecular phylogenetic analyses including *Histriobdella*, the genus presents a very long branch on the phylogenetic tree and tends to group with other long-branched lineages (Tilic et al., 2022). The relationships of such long-branched taxa are notoriously difficult to infer accurately on phylogenetic trees due to the long branch attraction artifact (Bergsten 2005; Kapli et al. 2020). When restricting its position to Eunicida, Tilic et al. (2022) found support for Histriobdellidae being closely related to the dorvilleid species included in their phylogenomic analysis, yet, this also presented a very long branch.

In the present study, we attempt to place Histriobdellidae within the Eunicida tree based on comprehensive taxon sampling of 52 taxa and a combination of two mitochondrial and two nuclear markers with high sequence data coverage. We furthermore reinvestigate the morphology of *Histriobdella homari* via Scanning electron microscopy (SEM) as well as based on standard immunohistochemical stainings and subsequent imaging with confocal laser scanning microscopy (clsm). Using such a comparative approach, we propose hypotheses on the putative homology of the head and body appendages between histriobdellids and other eunicidans. The evolution of Histriobdellidae’s aberrant morphology and suggested homologies are discussed relative to its phylogenetic position and commensal life style.

## Material and Methods

### Taxa selected for phylogenetic analyses

Fifty-two species representing six Eunicida families were used for the reconstruction of the phylogenetic position of Histriobdellidae within Eunicida (suppl. Fig. 2). New sequences were obtained from ethanol stored specimens of *Histriobdella homari*, *Stratiodrilus aeglaphilus*, two species of Eunicidae, three species of Dorvilleidae, five species of Lumbrineridae, two species of Oenonidae, and two species of Onuphidae (suppl. Fig. 2). Paragenophore vouchers of the newly sequenced species including *Histriobdella homari*, *Stratiodrilus aeglaphilus*, and hologenophore vouchers of 27 species of other enicidans were deposited at the collections of the University Museum of Bergen (ZMBN), the Australian Museum (AM) and the P.P. Shirshov Institute of Oceanology, Russian Academy of Sciences (IORAS) (suppl. Fig. 2). Data of 46 taxa were obtained from NCBI and BOLD (suppl. Fig. 2). The following taxa were not included in the analysis based on the previous knowledge of their extremely long branches on Eunicidan phylogenetic trees: *Ophryotrocha labronica* La Greca & Bacci, 1962 (Struck et al., 2006), *Oenone fulgida* (Lamarck, 1818) (Struck et al, 2006; Zanol et al. 2010) to mitigate the long branch attraction artifact in the phylogenetic analyses.

### DNA extraction and PCR

Genomic DNA was extracted from 96% Ethanol fixed samples using QuickExtract™ DNA Extraction Solution. One hundred µl of the solution was added to each air-dried tissue sample, incubated for 45 min at 65°C following by 2 min at 98°C. Amplification of the targeted regions of COI, 16S rDNA, 18 S rDNA, and 28S rDNA was performed with TaKaRa Ex Taq HS kit in a 25 μl reaction consisting of 1 µl of DNA template, 17.35 µl of purified water, 2.5 µl of 10x Ex Taq buffer, 2 µl of dNTP mixture, 1 µl of each primer and 0.15 µl of TaKaRa Ex Taq HS. Amplification protocols and primers used for each marker are shown in Table XX. The amplified products were purified and bidirectionally sequenced by Macrogen Europe (Amsterdam, Netherlands). Assembly and editing of raw sequence reads were done in Sequencher v. 4.5 (Gene Codes, Ann Arbor, Michigan). GenBank accession numbers and BOLD process IDs of all obtained sequences are listed in suppl. Fig. 2.

### Phylogenetic analyses

All sequence analyses were done on the CIPRES Science Gateway V. 3.3 except concatenation of the four genes into a single matrix that was done manually in MEGA 7 (Kumar et al. 2015).

#### Sequence alignment and masking

Sequences of individual markers were aligned using MAFFT (Katoh & Standley 2013) with default settings. The alignment with COI sequences was translated into amino acid sequences in Mega 7 using the invertebrate mitochondrial code (NCBI translation code 5) to ensure stop codons or frameshift mutations were not present. The sequences were then back translated into nucleotides for further phylogenetic analyses. To mitigate the long-branch attraction artifact, the aligned rRNA gene sequences were masked in Gblocks (Castresana 2000) to eliminate poorly aligned positions such as the long indels in the sequences of Oenonidae and Dorvilleidae. The following settings were used: maximum number of contiguous nonconserved positions – 8; minimum length of a block – 10; allowed gap positions – With Half.

#### Bayesian inference

Best-fit partitions and models were inferred with PartitionFinder2 on XSEDE v1.6.10. The selected models were: SYM+I+G for the 1^st^ codon position in COI; GTR+I+G for the 2^nd^ codon position in COI, 16S, and 18S; and GTR+G for the 3^rd^ codon position in COI and 28S. The PartitionFinder2 output file with the best partition scheme was used to compile the input Nexus file for the phylogenetic analysis in MrBayes v. 3.2.1 (Ronquist et al. 2012). Two independent and simultaneous runs with flat prior probabilities and four chains were run for 50,000,000 generations. Trees were sampled every 1000th generation. Stationarity of each chain was checked with TRACER v.1.8 (Rambaut et al, 2018) and the first 25% discarded as burn-in after visualizing a plot of likelihood score. The remaining trees were summarized into a majority rule consensus tree with posterior probabilities (PP) indicating the support value for each clade. Convergence between the runs was verified using the Average Standard Deviation of Split Frequencies (ASDSF) calculated in MrBayes. Tracer v. 1.8 (Rambaut et al. 2018) was used to examine MCMC sampling statistics and parameter estimates and to verify stationarity with plots of log likelihoods. An effective sample size (ESS) higher than 2000 for the log likelihood and all other parameters when the two runs were combined was considered a good mixing, and the results of the analyses were accepted.

#### Maximum likelihood

Maximum Likelihood analysis for the combined dataset was conducted using IQ-TREE multicore version 2.1.2 (Nguyen et al. 2015) with 5,000 bootstrap (BS) replicates, 300 initial parsimony trees, 15 trees to be maintained in the candidate set during ML tree search and the automatic model selection option. The models selected during the analysis were: SYM+I+G4 for the 1^st^ codon position in COI, K3Pu+F+I+G4 for the 2^nd^ codon position in COI, TN+F+G4 for the 3^rd^ codon position in COI, TIM2+F+I+G4 for 16S, TIMe+I+G4 for 18S, and TIM3+F+G4 for 28S.

### Morphological studies

#### Sampling and fixation of Histriobdellidae specimens

Specimens of *Histriobdella homari* Beneden, 1858 were collected from their host, the European lobster *Homarus gammarus* (Linnaeus, 1758) (Nephropidae, Crustacea) purchased in December 2016 and June 2017 at Bergen fish market, presumably collected locally in Norway.

To detach the histriobdellids from their host, the respective parts of the lobster were washed using 7% MgCl_2_ in FSW. Specimens for subsequent DNA extraction were stored in 96% ethanol. Specimens for immunohistochemistry studies were fixed in 4% paraformaldehyde (PFA) in 1x phosphate buffered saline containing Tween (PTW: 0.05 M PB/ 0.3 M NaCl/ 0.1% Tween20) for 2h at room temperature (RT). Afterwards, histriobdellids were rinsed in PTW several times and stored in PTW containing 0.05% NaN_3_ at 4°C until usage. For electron microscopy, animals were fixed in 2.5% glutaraldehyde in sodium cacodylate buffer (0.1 M Cacodylate, pH 7.4, 0.24 M NaCl) for 1h at RT and stored at 4°C in sodium cacodylate buffer until usage.

#### Scanning electron microscopy (SEM)

For SEM analyses, 15 fixed specimens of *H. homari* were washed 2x5 min in sodium cacodylate buffer (0.1 M Cacodylate, pH 7.4, 0.24 M NaCl). Afterwards, the specimens were postfixed in osmium tetroxide (1% O_S_O_4_ in sodium cacodylate buffer) for 45 min at RT and rinsed with distilled water for 3x10 min. Subsequently, the samples were dehydrated using an increasing EtOH series (30%-40%-50%-60%-70%-80%-90%-95%-3x100%, 5-10 min each), critical-point dried and coated with gold/palladium.

Finally, the samples were investigated with a Supra 55VP scanning electron microscope (Zeiss, Germany). The final panels were designed using Adobe (San Jose, CA, USA) Photoshop CC and Illustrator CC.

#### Immunohistochemistry and confocal laser scanning microscopy (clsm)

Antibody stainings using standard markers were performed on whole animal preparations. Although the specificities of the used antibodies have all been established in numerous invertebrates (for references see discussion), we cannot exclude that a given antibody may bind to another related antigen in the investigated specimens. Therefore, we refer to observed signals as exhibiting (antigen-) like immunoreactivity (-LIR). Due to the specificity of the anti-tubulin staining, we refer to (antigen-) immunoreactivity (-IR) in this case. Negative controls were conducted by omitting the primary antibody in order to check for antibody specificity and yielded no fluorescence signal.

At least 20-30 adult specimens of *H. homari* using the cslm. PFA-Fixed and stored individuals were rinsed 2x5 min in PTW (PBS with 0.1% Tween 20) at room temperature (RT) and subsequently transferred into 10 µg proteinase K/ml PTW for 10-15 min to penetrate the membranes. After two short rinses in glycine (2 mg glycine/ml PTW), and 3x5 min washes in PTW, specimens were fixed a second time using 4% PFA in PBS containing 0.1% Tween for 20 min at RT, rinsed 2x5 min in PTW, 2x5 min in THT (0.1M Tris-HCl pH 8.5, 0.1% Tween-20) and blocked for 1-2 h in 5% sheep serum in THT. The primary antibodies, polyclonal rabbit anti-CFLRFamide (GenScript, Piscataway, USA, dilution 1:250) and monoclonal mouse anti-acetylated α-tubulin (Sigma-Aldrich, St. Louis, USA, dilution 1:250), were applied for 48-72 h in THT containing 5% sheep serum at 4°C. Afterwards, specimens were rinsed twice in 1 M NaCl in THT, then washed 5x30 min in THT and incubated subsequently with secondary fluorochrome conjugated antibodies (goat anti-rabbit Alexa Fluor 488, Invitrogen, USA, dilution 1:500; goat anti-mouse Alexa Fluor 633, ANASPEC, Fremont, USA, dilution 1:500) in THT containing 5% sheep serum for 48 h at 4°C. Subsequently, samples were washed 6x30 min in THT, stained with DAPI for 15-30 min (5mg/ml stock solution, working solution: 2µl in 1 ml THT – final concentration 10µg/ml) and/or phalloidin-rhodamine (5 µl methanolic stock solution in 500 µl THT) and washed 2x5 min in THT.

For anti-synapsin staining, specimens were fixed overnight in 1 % aqueous formaldehyde containing ZnCl_2_ (18.4 mM), NaCl (135 mM) and sucrose (35 mM)). After several rinses in Hepes-buffered saline (HBS: 10 mM HEPES, 25 mM saccharose, 150 mM NaCl, 5 mM KCl, 5 mM CaCl_2_) for at least 3 × 15 min, specimens were incubated in an 80 % methanol/20 % DMSO mixture for 1.5 h at RT. Subsequently, specimens were transferred to 100 % methanol for at least 1 h at RT. Afterward, the samples were rehydrated stepwise and finally incubated in Tris-buffered saline (TBS: 50 mM Tris, 150 mM NaCl, pH 7.4, 10 min, “buffer I”). Following this, the samples were transferred into TBS containing 5 % normal goat serum (NGS, Sigma-Aldrich, St. Louis, MO, USA, 1 % DMSO and 0.0005 % NaN_3_: “buffer II” for 1 h at RT). This was followed by incubation in the primary antiserum (1:30 in buffer II) for 72 h at 4 °C. Afterward, specimens were rinsed in buffer II for 3 × 2 h and subsequently incubated in secondary antibody (goat anti-mouse Alexa Fluor 568, ANASPEC, Fremont, USA, 1:1,000 in buffer II, 48 h, 4 °C). Thereafter, the specimens were washed once in buffer II and twice in buffer I.

Finally, incubated specimens were then dehydrated using an ascending isopropyl alcohol series, transferred into Murray’s Clearing solution (2 parts benzyl benzoate and 1 part benzyl alcohol) and subsequently mounted between two cover slips using DPX slide mounting medium (Sigma-Aldrich, St. Louis, USA). Specimens were analysed with the confocal laser-scanning microscope Leica TCS SP5 (Leica Microsystems, Wetzlar, Germany). Confocal image stacks were processed with Leica AS AF v 2.3.5 (Leica Microsystems) and Imaris 9.3 (Bitplane AG, Zurich, Switzerland). The final panels were designed using Adobe (San Jose, CA, USA) Photoshop CC and Illustrator CC.

## Results

### Molecular data

The combined data set has 4268 aligned positions (COI with 659 positions, mitochondrial 16S rRNA with 576 position 18S rRNA with 2141 positions, and 28S rRNA with 892 positions). After applying Gblocks the new 16S rDNA alignment retained 313 positions (54%), 18S rRNA alignment retained 1582 positions (74%), 28S rRNA alignment retained 511 positions (57%). Tree topologies obtained in the Bayesian and ML analyses were highly congruent. All eunicidan families were recovered as monophyletic with high support: Dorvilleidae (PP=1.00, BP=92), Eunicidae (PP=0.99, BP=91), Histiobdellidae (PP=1.00, BP=100), Lumbrineridae (PP=1.00, BP=100), Oenonidae (PP=1.00, BP=100), Onuphidae (PP=1.00, BP=95). Eunicidae was sister to Onuphidae (PP=1.00, BP=100) and Dorvilleidae was sister to Histriobdellidae (PP=1.00, BP=92). Lumbrineridae were sister to all other families included into the analysis (PP=1.00, BP=79). On the Bayesian tree, Onuphidae+Eunicidae, Histiobdellidae+Dorvilleidae and Oenonoidae formed a polytomy (Fig. 1), while on the ML tree, Oenonidae were sister to (Onuphidae+Euniciae, Histriobdellidae+Dorvilleidae) although with poor support (BP=36) (suppl. Fig. 1). Although investigation of the relationships within each family clade was beyond the scope of the present study, the following key findings can be outlined. *Lumbrineris*, the type species of the family Lumbrineridae appeared polyphyletic. Within Eunicidae, the genera *Marphysa* (PP=0.97, BP=66), *Paucibranchia* (PP=1.00, BP=100) and *Palola* (PP=0.99, BP=80) were recovered as monophyletic, while *Eunice* and *Leodice* were paraphyletic. Within Onuphidae two sister clades were recovered, representing the two subfamilies: Hyalinoeciiae (PP=1.00, BP=100; with *Hyalinoecia*, *Leptoecia,* and *Nothria*) and Onuphinae (PP=0.86, BP=45; with *Austarlonuphis*, *Diopatra*, *Hirsutonuphis*, *Mooreonuphis*, *Onuphis*, *Paradiopatra*, *Paxtonia*, and *Rhamphobrachium*). Within Oenonidae, *Tainokia* was sister to *Arabella*+*Drilonereis* (PP=1.00, BP=100). Two well supported sister clades were recovered within Dorvilleidae: the first clade included *Dorvillea*, *Protodorvillea* and *Schistomeringos* (PP=1.00, BP=100) and the second clade comprised *Ougia* and *Parougia* (PP=1.00, BP=100).

**Fig. 1:**
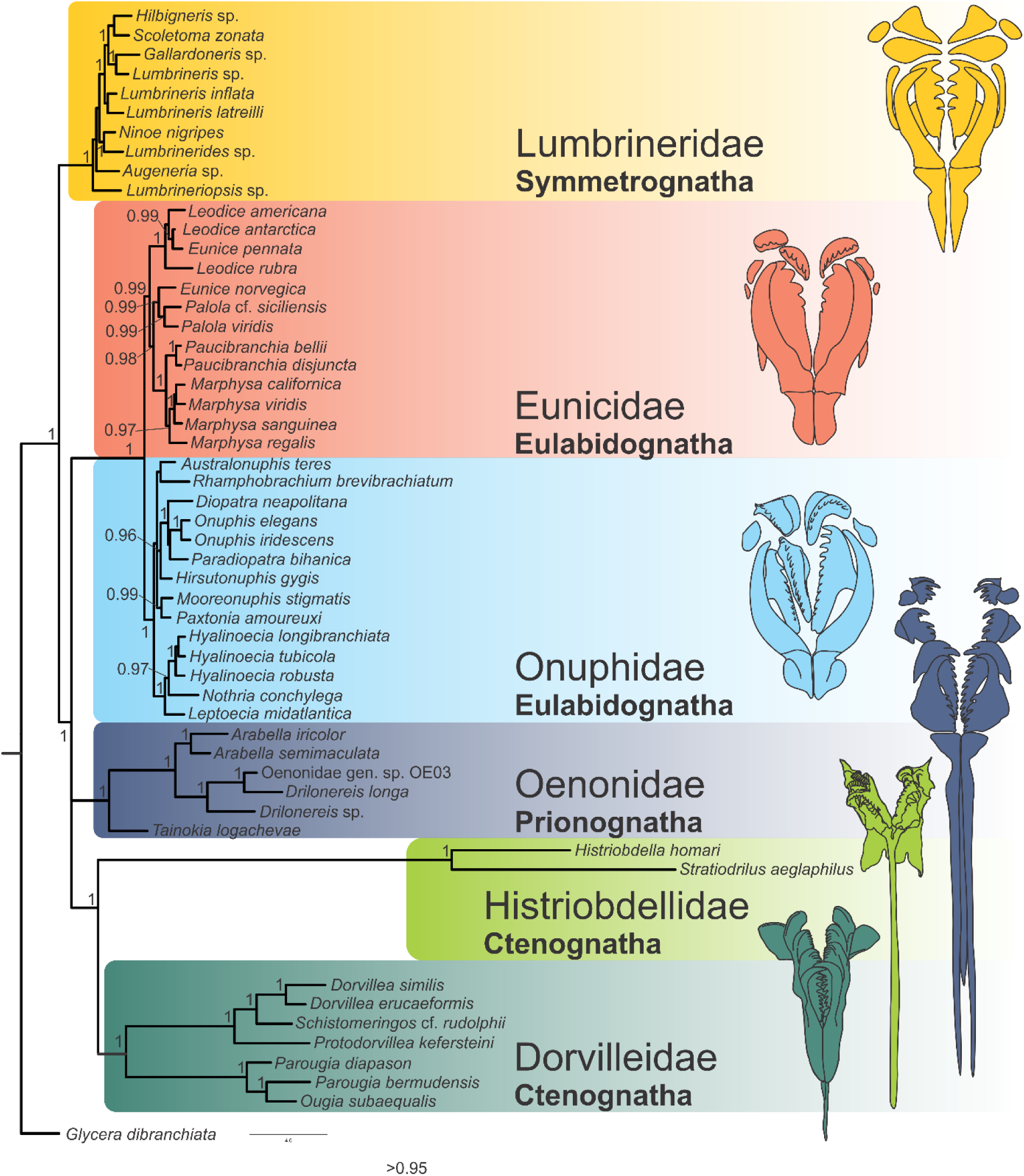
Best Baysian Inference tree resulting from a combined data set including COI, 16S rDNA, 18S rDNA and 28S rDNA. The tree topology include 53 taxa out of almost all families within the Eunicida (except Hartmaniellidae). The colour-coded drawings illustrate the shape of the jaw apparatuses in the respective families.

### Morphological investigations

#### The general morphology of adult *Histriobdella*

As already described by various authors, adult specimens of *Histriobdella homari* exhibit a minute, body with at least six externally recognisable, elongated segments (Fig. 2a). They show a distinct anterior end bearing five putatively sensory appendages as well as one pair of so-called adhesive papillae, whereas the posterior end is characterized by one pair of lateral foot-like appendages (Fig. 2a). The internal anatomy shows a prominent brain neuropil and at least eight ganglia along the trunk (Fig. 2b). Nevertheless, further putative ganglia are present within the posterior appendages according to anti-synapsin staining (Fig. 2b). Furthermore, the nervous system exhibits a distinct ventral nerve cord as well as innervating neurite bundles leading towards all appendages (Fig. 2d). The musculature is characterized by the lack of circular muscle bundles. Instead, numerous longitudinal muscles run along the body (Fig. 2c). In the following, the different appendages will be characterized in detail and main focus will be given to their neuronal and muscular innervation.

**Fig. 2:**
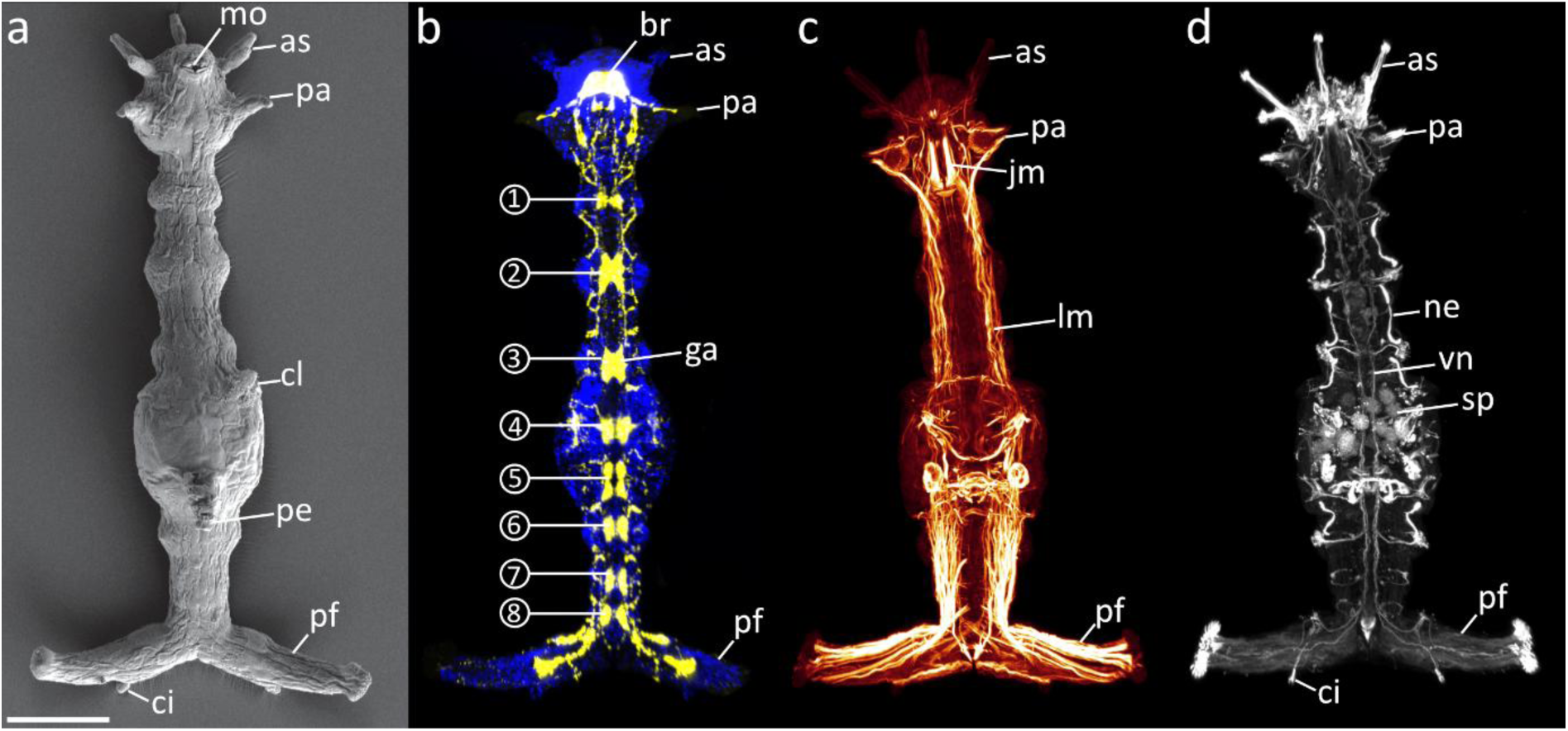
External and internal morphology of *Histriobdella homari*, modified from Helm, Vila & Budeava (2020). a. SEM image, b-c. confocal maximum projections. Anterior is up in all images and all specimens are shown in ventral view. a. The adult morphology is characterized by five prominent anterior sensory appendages (as), so-called adhesive papillae (pa) and distinct posterior foot-like appendages (pf). B. Anti-synapsin staining reveals presence of at least eight main ganglia (ga, 1-8) along the trunk. Notably, additional neuropils are visible in the foot-like appendage (pf). C. The musculature of adult specimens is characterized by distinct longitudinal muscle bundles (lm), but the lack of circular musculature. d. Anti-tubulin staining exhibits a strong neuronal innervation of the anterior appendages (as, pa) as well as a distinct ventral nerve cord (vn). as, anterior sensory appendage; br, brain; ci, cirrus-like structure; cl, clasper; ga, ganglion; jm, jaw musculature; lm, longitudinal musculature; mo, mouth opening; ne, nephridia; pa, adhesive papillae; pe, penis; pf, posterior foot-like appendage; sp, sperm; vn, ventral nerve cord; 1-8, number of ganglia along the trunk. Scale bar = 100 µm

#### The anterior-most sensory appendages

In *Histriobdella homari*, the prostomium bears five prominent and highly similar anterior appendages. Following the nomenclature of other eunicidans the dorsally-located three inner ones (Fig. 3a, II+III) are called antennae whereas the pair of ventro-lateral oriented structures are called palps (Fig. 3a, I). SEM reveals prominent ciliation on all of the five anterior-most appendages, each with a distinct group of longer cilia at the tip and an area of shorter cilia at the ventrally-oriented proximal part (Fig. 3a, inset). A staining against acetylated α-tubulin exhibits the presence of dense neurite bundles along the proximal part of each appendage (Fig. 3b). In each appendage a proximally-located assemblage of neuronal somata is present. According to the structure and density of the somata, these will hereafter be referred to as appendage ganglia (Fig. 3c). These prominent structures – formerly described as brain lobes by Gelder and Jennings (1975) – innervate the anterior appendages, with the apical (putatively sensory) cilia, and connect the anterior appendages with the brain neuropil (Fig. 3d). The configuration of appendage nerves is somewhat similar although the number of bundles and their connection to the brain may differ (Fig. 3c, d). Thus, the median-most anterior appendage (Fig. 3d, III) comprises several distinct neurite bundles connected to the dorsal root of the circumesophageal connective (see also Fig. 6a for schematic overview). The dorso-lateral pair of appendages (Fig. 3d, II) each contains only two neurite bundles solely connected to the dorsal root of the circumesophageal connective (see also Fig. 6a for schematic overview). Contrary, the two neurite bundles of the ventro-lateral pair of appendages (Fig. 3d, I) show a distinct connection to both the dorsal and ventral root of the circumesophageal connective (see also Fig. 6a for schematic overview). Clsm images of F-actin stained muscles show a strong and comparable muscular innervation of the base of each anterior appendage (Fig. 3e). Notably, muscle bundles are almost absent within each appendage. The innervation is similar between the laterally-oriented two pairs of appendages with each appendage being supported by minimum three distinct muscles, whereas the median-most appendage is supported by fewer F-actin fibres (Fig. 3e).

**Fig. 3:**
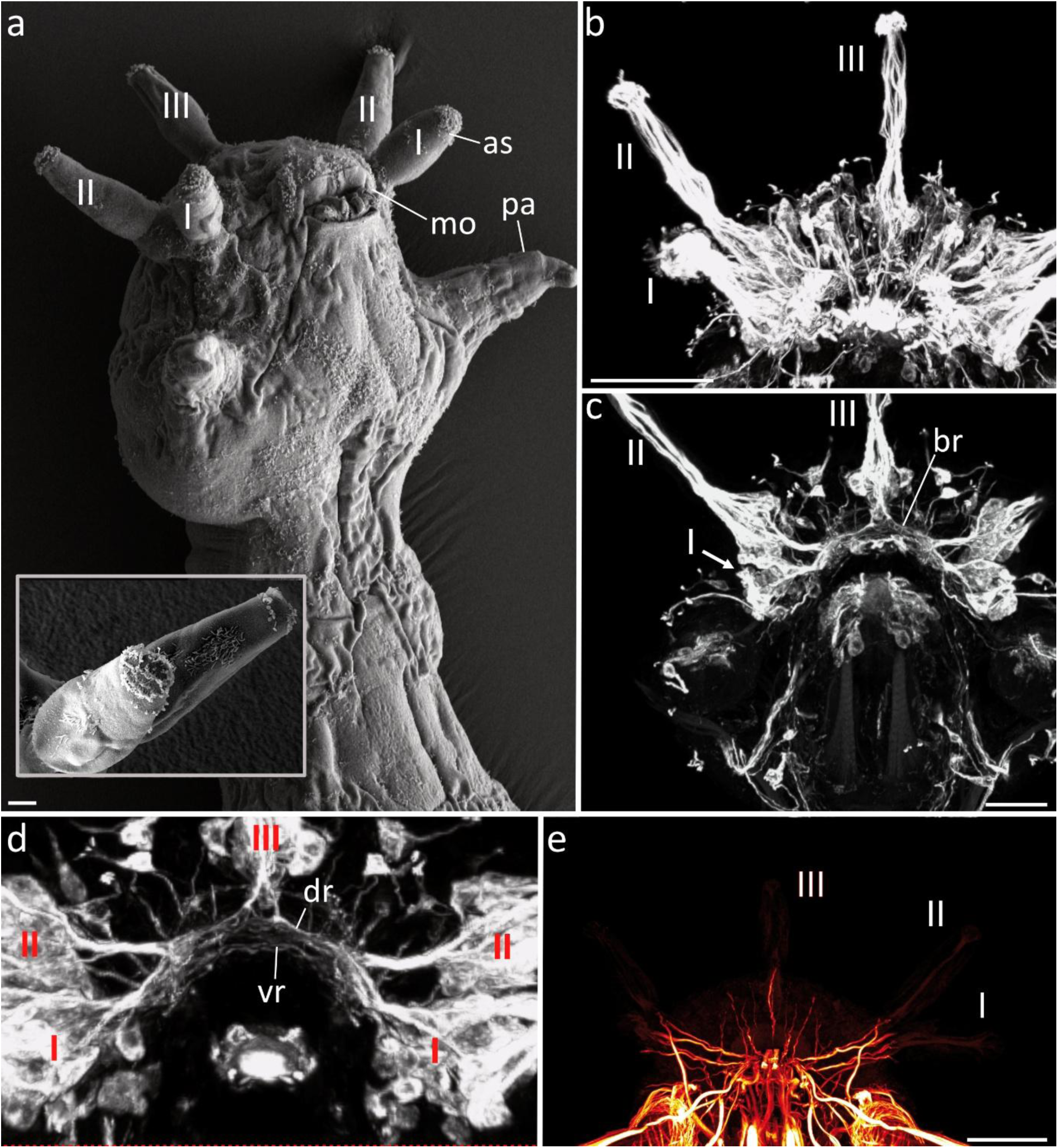
The anterior sensory appendages observed via SEM (a) and clsm (b-e). a. The anterior sensory appendages (as, I-III) exhibit a dense ciliation at the tip and the ventral shaft of the structure. The inset shows a higher magnification of I and II in anterior view. b, c. Anti-tubulin staining reveals assemblages of somata at the base of each anterior appendage and all are closely connected with the brain region (br). c. The median (III) and lateral (II) anterior appendages are innervated by neurite bundles coming from the dorsal root of the brain (dr), whereas the neurite bundles innervating the ventro-lateral anterior appendages (as) originate in the dorsal (dr) and ventral root (vr). e. Although the neuronal innervation differs, all anterior appendages (as, I-III) show a similar pattern of muscular innervation. as, anterior sensory appendage; br, brain; dr, dorsal root of the circumesophageal connective; pa, adhesive papillae; ventral root of the circumesophageal connective. Scale bars = 10 µm (a,), 30 µm (b-e).

#### Structure and innervation of the adhesive anterior papillae

The adhesive anterior papillae are conical outgrowths of the body wall with a distal swelling (Fig. 4a). The proximal part is filled by a dense bundle of glands which long necks extend to the tip and open separately in a row of pores (Fig. 4d-f). A row of single cilia is found along the posterior side of the tip (Fig. 4e). Anti-α-tubulin-IR shows two distinct neurite bundles running from proximal towards the distal end of the papillae. Furthermore, a peripheral neurite bundle branches off along the way and forms an extra neurite reaching the distal end. These neurite bundles represent a continuation originating at the circumesophageal connectives and interconnect the entire structure with the latter (Fig. 4a, 6b). Additionally, distinct α-tubulin-IR is present in the walls of the glandular necks. Anti-CFLRF-amide staining also shows a strong innervation of the papillae via two neurite bundles running from the circumesophageal connective towards the apex (Fig. 4b). Staining against synapsin exhibits a dense neuropil within the appendage, leading from the basal parts towards the tip. Furthermore, the prominent interconnection of the appendage with the circumesophageal connective close to the brain is shown via synapsin-LIR (Fig. 4c).

**Fig. 4:**
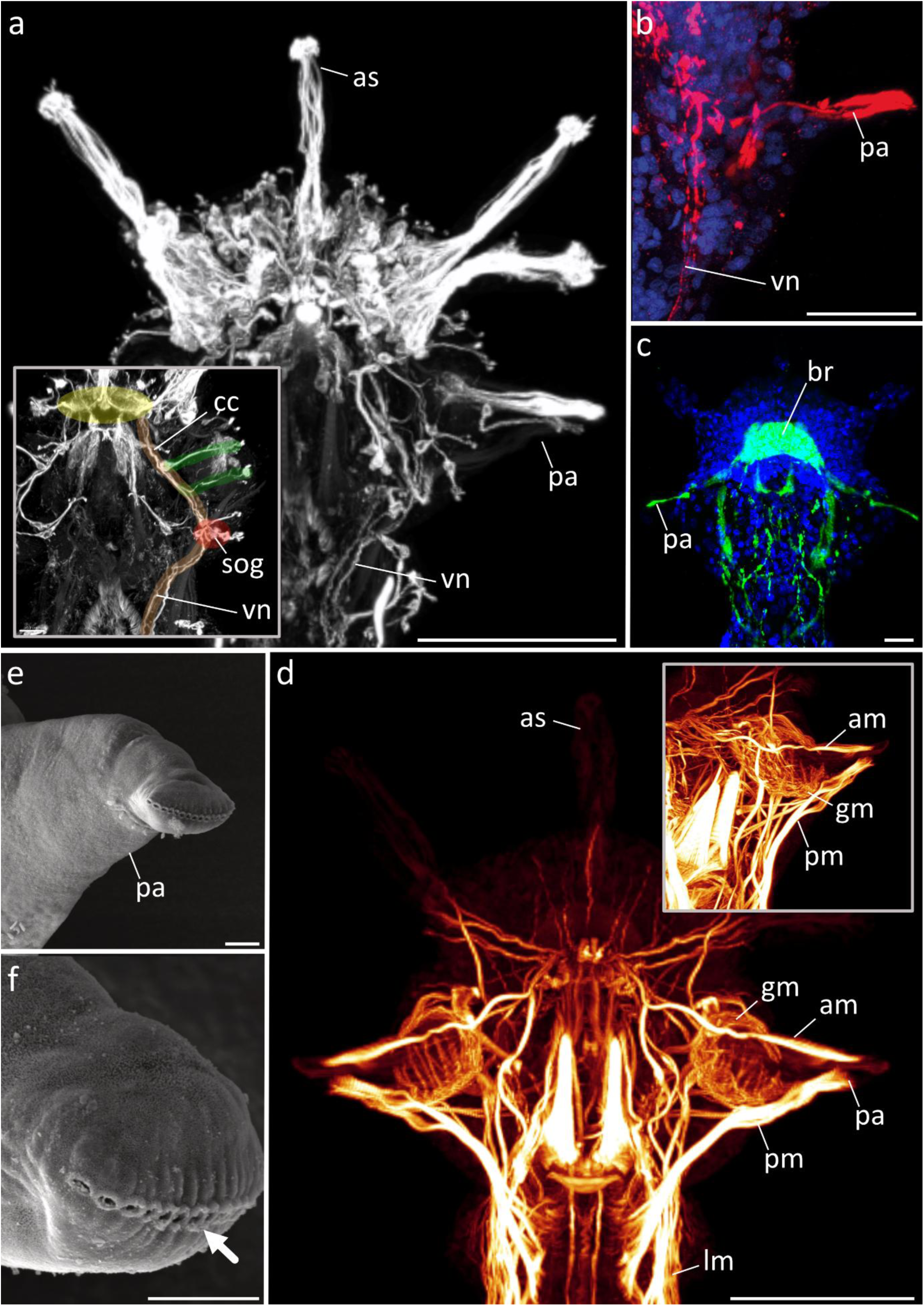
The anterior adhesive papillae shown via confocal maximum projections (a-d) and SEM (e, f). a. Anti-tubulin staining reveals a strong neuronal innervation. The inset shows the color-coded neuronal scaffold: brain (yellow), elongation of the ventral nerve cord (vn) leading into the brain (orange), neurite bundles running into the adhesive papillae (green) and the position of the subesophageal ganglion (sog) (red) posterior of the latter. b. Staining against CFLRF-amide highlights the presence of an innervating neurite bundle running into the structure and originating from the ventral nerve cord (vn). c. Anti-synapsin staining shows a prominent neuropile within the appendage (pa). d. Muscular staining exhibits distinct anterior (am) and posterior muscle bundles (pm) inside the papillae. A dorsal view (inset) reveals the presence of dorsal and ventral bundles forming the main muscles (am, pm) as well as the lack of additional muscle fibres. e, f. SEM shows distinct pores (arrow) at the tip of the papillae (pa). am, anterior muscle bundle; as, anterior sensory appendage; br, brain; cc, circumesophageal connective; gm, glandular muscle; lm, longitudinal muscle bundles; pa, adhesive papilla; pm, posterior muscle bundle; sop, subesophageal ganglion; vnc, ventral nerve cord. Scale bars = 50 µm (a, d), 30 µm (b, c), 2 µm (e, f),

Another interesting fact is the position of the subesophageal ganglion which is located halfway along the pharynx region – showing anti-tubulin-LIR and anti-synapsin-LIR (Figs. 4a inset, 4c, 6b) – and shows two widely separated ganglia situated along the two branches of the ventral nerve cord that run in the direction of the brain. Notably, the neurite bundles innervating the papillae branch off anterior of the respective ganglia (Fig. 4a, inset). F-actin staining shows the presence of dense muscle bundles within the papillae, not belonging to the body wall musculature (Fig. 4d). Instead, these muscle bundles run from the apical end of the structure towards its base, fan out there and root the entire appendage within the body wall muscles (Fig. 4d). Additionally, the anterior appendage muscles are interwoven into the musculature of the anterior end, whereas the posterior appendage muscles insert ventral of the longitudinal body wall muscles. Besides the anterior and posterior main muscle bundles, distinct musculature covering the ventral or dorsal part of the appendage is lacking (Fig. 4d, inset). Nevertheless, the anterior and posterior main bundles are each composed of dorsal and ventral muscles forming the respective main bundles within the appendage (Fig. 4d, inset). Such a muscular scaffold supports a putative function as retractive muscles, involved in releasing the grip of the papillae when detaching. Furthermore, F-actin staining reveals well-developed glandular musculature surrounding the glandular cluster proximally, and thinner muscles extending apically halfway along the gland necks, possibly controlling their secretory release (Fig. 4d).

#### The posterior appendages

The posterior-most appendages of *Histriobdella* are paired elongated foot-like structures exhibiting pore-like glandular openings at the distal end (Fig. 5a). Furthermore, a bulb-shaped cirrus is present halfway along the “feet” (Fig. 5a, b). According to our observations, the latter cirrus shows distinct apical ciliation supporting a putatively sensory function of the structure (Fig. 5a, insert). When observing the prominent neuronal innervation, this function seems plausible (Fig. 5c).

**Fig. 5:**
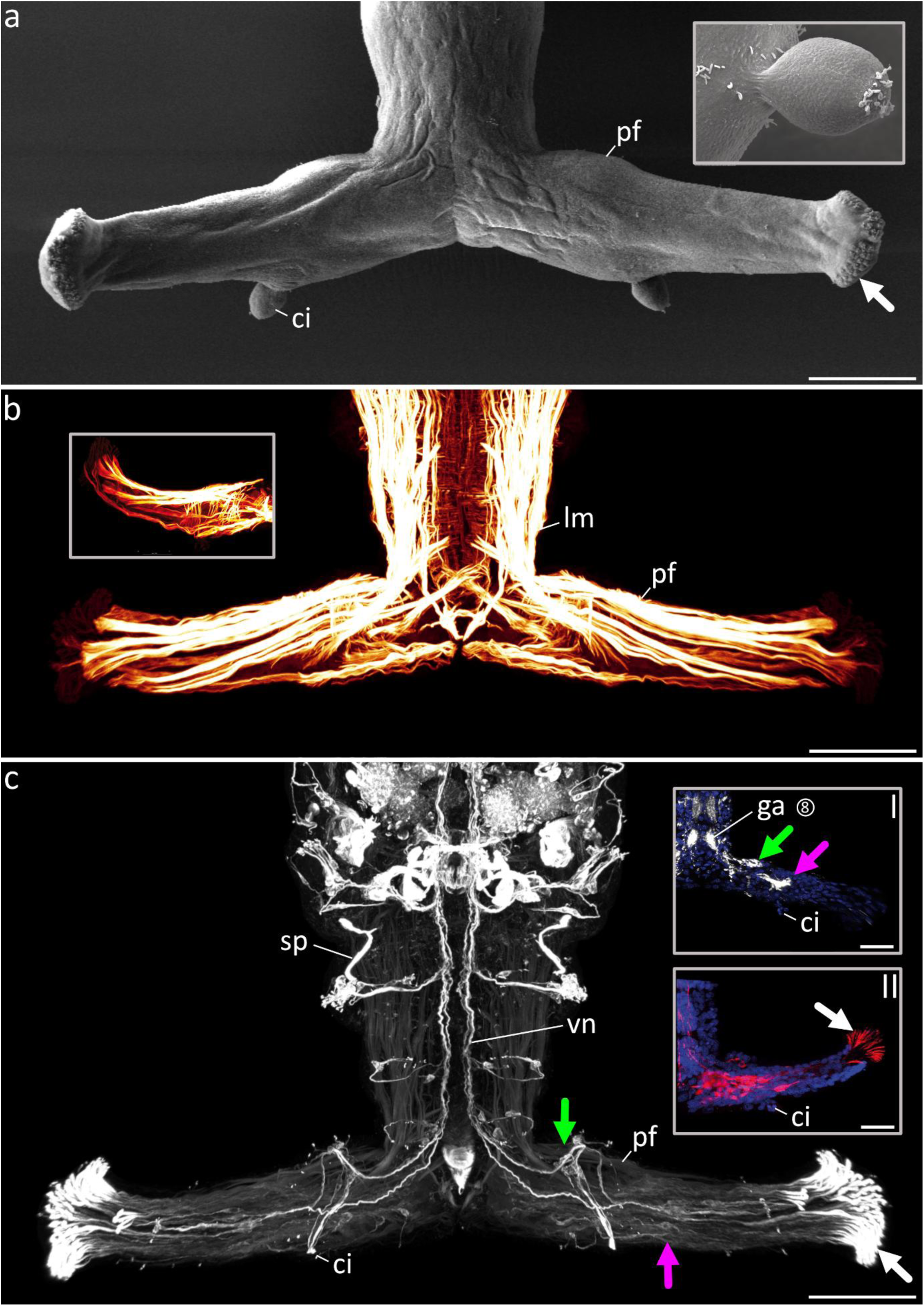
The posterior foot-like appendage shown via SEM (a) and confocal maximum projections (b, c). a. The posterior foot-like appendages are characterized by apical glandular openings (arrow) and a cirrus-like structure (ci) attached posteriorly (see also inset for higher magnification). b. b. The musculature of the posterior end of *Histriobdella* is characterized by distinct longitudinal muscles (lm) of the body wall extending into the posterior foot-like appendages (pf). Furthermore, prominent bracing muscles are present (inset). d. Anti-tubulin staining reveals the presence of distinct neurite bundles representing a continuation of the ventral nerve cord (vn). The cirrus-like appendage (ci) is innervated by neurites coming from a ganglion situated along a anterior subset of neurite bundles of the *vn* (green arrow). Afterwards all neurite bundles within the *pf* run into a second ganglion (pink arrow) before entering the glandular structures (white arrow). Anti-synapsin staining supports the presence of both ganglia. Anti-CLRF-amide staining also exhibits prominently marked somata along the *pf*. ci, cirrus-like appendage; ga, ganglion; pf, posterior foot-like appendage; sp, sperm; vn, ventral nerve cord. Scale bars = 50 µm (a-c).

Along the entire posterior appendage, two pairs of neurite bundles representing an elongation of the histriobdellid ventral nerve cord and originating from ganglion 8 extend into the structure (Fig. 5c, insert II). The anterior neurite bundle of each appendage bears an additional ganglion at the proximity of the appendage (putative ganglion 9; Fig. 5c, green arrow), which gives rise to neurites innervating the cirrus (Fig. 5c, 6c). Posterior to the latter ganglion, both main neurite bundles of each appendage terminate in another additional ganglion at the distal end of the structure, close to the distal glandular openings (Fig. 5c, pink arrow). The long gland necks show strong α-tubulin staining (Fig. 5c) caused by a tubulin-rich outline of the glandular pores (see discussion). Furthermore, anti-synapsin staining of the posterior appendage verifies the presence of the prominent ganglia within the appendage (green and pink arrows, Fig. 5c, insert I). Anti-CFLRF staining reveals immunoreactivity within distinct cell clusters within the region of the two ganglia as well (Fig. 5c, insert II). In case of the anti-CFLRF staining, the positive signal within the gland channels leading towards the glandular openings has to be considered as being a false positive signal (Fig. 5c, insert II, white arrow).

F-actin staining shows a strong muscular innervation of the posterior paired appendages (Fig. 5b). Hence, the longitudinal body wall musculature of the trunk runs into the appendages and forms major parts of the muscular scaffold (Fig. 5b). Nevertheless, few appendage-specific sets of muscle fibres are present, including dense bracing muscles underlying the longitudinal ones at the proximity of the appendage (Fig. 5b, inset). Furthermore, the distal-most muscle bundles do not represent an elongation of the body wall fibres, but are formed by appendage-exclusive F-actin bundles (Fig. 5b).

## Discussion

### Phylogenetic position of Histriobdellidae

Despite the extremely modified body plan, histriobdellids were recognized as a member of Eunicida due to the presence of the complex jaw apparatus in the ventral muscular pharynx (Hatschek 1888, Mesnil and Caullery 1922, Jennings and Gelder 1976, Fauchald and Rouse 1997, Paxton 2009). Nevertheless, the position of histriobdellids within Eunicida has been debated. Tzetlin et al. (2020) suggested the close relationships between Histriobdellidae and Dorvilleidae based on the similarity in the ultrastructure of the maxillary plates. In phylogenetic analyses based on the transcriptomic data Tilic et al. (2022) reported conflicting hypotheses on the placement of Histriobdellidae, either as sister to Dorvilleidae when restricting their position to within Eunicida or in various poorly supported positions outside Eunicida in non-restricted analyses. However, their analysis was based on a single representative from each eunicidan family except Hartmaniellidae (absent in the analysis). The conflicting topologies were explained by the possible long branch attraction artifact.

Histriobdellids are frequently described as aberrant commensal animals that are considered of problematic phylogenetic placement due to their very long branches on phylogenetic trees (Tilic et al. 2022). A variety of strategies are known to be used to alleviate the LBA artifact: increase in taxon coverage excluding long branches, removing unreliably aligned regions, excluding third codon positions; translating into amino acids in protein coding genes (see review in Bergsten 2005). In large genomic datasets; filtering data by removing rapidly evolving genes or sites (Kapli et al. 2020); applying the site-heterogeneous CAT model (Lartillot et al. 2007) or coalescent-based tree-building methods (Liu et al. 2015) can be also used.

In the present study we attempted to mitigate the long branch attraction artifact by increasing the taxon and sequence data coverage, excluding other known taxa with long branches from the analysis, and by removing large indels and poorly aligned regions from the highly divergent sequences applying Gblocks masking. Despite that our molecular dataset was based on only four molecular markers, the taxon and sequence data coverage for each family was the most comprehensive up to date with representatives of 33 eunicidan genera (33% of known total generic diversity) including two known histriobdellid genera: *Histrobdella* and *Stratiodrilus*. Both 18S rRNA and 16S rRNA had no missing sequence data, while only 13% and 36% of sequences were missing in COI and 28S rRNA respectively. Applying Gblocks to the three ribosomal RNA genes, resulted in eliminating nearly 25–50% of poorly aligned positions.

Although histriobdellids do show the longest branch on our phylogenetic tree, no other long-branched taxa were present. Strongly supported sister relationships between Histriobdellidae and Dorvilleidae corroborate earlier proposed hypotheses on their affinity (Tzetlin et al. 2020, Tilic et al. 2022) and suggest that their jaws are of the ctenognath type (Tzetlin et al. 2020). Ctenognath maxillae bear numerous very small denticles suggesting that they are used for microphagy – grazing on bacterial or unicellular algae, a feeding mode characteristic for both dorvilleids and histriobdellids (Rouse, Pleijel & Tilic, 2022). This similarity in feeding strategy also supports the putative sister relationships between minute histriobdellids and dorvilleids. In contrast, all other eunicidan annelids are characterized primarily as macrophagous feeders, i.e. predators, scavengers or herbivorous grazers (Rouse, Pleijel & Tilic, 2022) with maxillary apparatuses of different types (prionognath, eulabidognath or symmetrognath) but all bearing large denticles on their maxillary plates.

### The anterior-most appendages – palps or antennae?

According to our comprehensive datasets, the anterior-most appendages of *Histriobdella homari* are highly similar in terms of ciliation and their muscular and neuronal scaffold. Hence, all five appendages share the lack of distinct muscle bundles within the appendage itself, but show a dense and comparable meshwork of muscle fibres at the base of the structure. Furthermore, the assemblage of neurites within the appendages visualised with anti-acetylated α-tubulin IR are highly comparable (see also Gelder & Jennings, 1975). This similarity could not be corroborated by IR against anti-CFLRF-amide and anti-synapsin, lacking in the appendages. The only and most-obvious difference between all five anterior-most appendages is their position of origin in terms of innervating neurites branching off from the central nervous system. According to morphological investigations by Orrhage (1995) in histriobdellids, the three median anterior appendages should be called antennae whereas the ventro-lateral ones should be referred to as palps based on their neuronal innervation pattern. Same is true for the head appendages of other eunicid families (Zanol, 2010; Kuhl et al., 2022). Accordingly, palps in Eunicida (and most remaining Annelida) are characterized by a neuronal innervation originating in both roots of the circumesophageal connectives, whereas antennae are solely innervated by neurite bundles coming from the dorsal root. Our observations support such a pattern and therefore verify previous homology assumptions for Histriobdellidae as well. Nevertheless, the neuronal details exhibited by our investigations add knowledge to previous studies and build up a backbone for comparative analyses. Furthermore, the observed innervation patterns are in line with neuronal scaffolds present in other errant taxa, such as e.g., in Syllidae (Schmidbaur et al., 2020). When it comes to palps in general, similar neuronal innervation patterns are also known for non-errant Annelida (Orrhage & Müller 2005; Purschke 2016; Kalke et al., 2024).

### Parapodial affinities of the adhesive anterior papillae

The adhesive locomotory papillae possess a number of characters that make them highly comparable with annelid parapodia. The muscular arrangement with muscle fibres exclusively supporting the locomotory appendage and not branching off from the body wall musculature represents a parapodia-like feature (Tzetlin & Filippova, 2005). Furthermore, the prominent insertion of the appendage main muscles into the existing longitudinal body wall muscles represents a parapodia-like scaffold known for other errant annelids as well. Nevertheless, the histriobdellid condition that the dorsal part of the anterior and posterior appendage muscle bundles innervate ventral of the dorsal longitudinal body wall muscles, before inserting in the dorsal midline or rather intertwine with the dorsal longitudinal muscle bundles, mirrors the condition observable in most annelid taxa (see also Allentoft-Larsen et al, 2021). In the latter, the parapodial main muscles are attached dorso-transverse of the ventral body wall muscle bundle. Additionally, in *Histriobdella* the anterior appendage muscle bundle – as well as the posterior one – is actually composed of dorsal as well as ventral muscles. The latter condition is therefore similar to parapodial flexor, protractor and extensor muscles observable in other clades (e.g., Storch, 1968; Müller & Worsaae 2006; Winchell et al., 2010; Allentoft-Larsen et al. 2021). In fact, the complexity of the muscular scaffold differs from that of most other annelid groups in the number of involved muscle fibres, which seem quite reduced in *Histriobdella*. Such a limited number of parapodial muscular elements might be the result of the mainly adhesive function of the locomotory anterior appendages in histriobdellids and the lack of additional structures such as acicula or chaetae otherwise supported by separate muscle bundles (e.g., Mettam 1967; Storch 1968; Filippova et al. 2006; Allentoft-Larsen et al. 2021). Comparable changes or reductions of the parapodial muscular arrangement due to lifestyle adaptations or the presence of solely uniramous parapodial structures are also known for other annelid families, such as e.g., the commensal Myzostomida (Helm et al., 2013), swimming scale worms (Allentoft-Larsen et al. 2021), or the benthic Sphaerodoridae (Helm & Capa, 2015).

When observing the neuronal innervation of the adhesive papillae, the picture gets more complex. Hence, the histriobdellid neurite bundles innervating the papillae branch off anterior to the so-called subesophageal ganglion and run towards the appendage tip (Gelder and Jennings, 1975). The fact, that these subesophageal ganglia are not situated posterior of, but halfway along the pharynx, supports a hypothesis that the position of the ganglia and the closely related papillae is the result of an evolutionary translocation via cephalization. Such a cephalization during evolution and/or development is known for several annelid groups, such as Nereididae (Fischer et al, 2010), Tomopteridae (Akesson, 1962; Purschke & Helm, 2023), Terebelliformia (Kalke et al., 2021), or Nerillidae (Worsaae, 2005). Thus, the first parapodial segment(s) is (are) integrated into the formation of the anterior end. As a result, the respective ganglia and parapodial structures are translocated towards anterior and often show a different shape (or are widely separated, such as in the case of the ganglia) when compared with the remaining parapodial segments. In Nerillidae, these translocated segments are called buccal segments (Worsaae, 2005). According to our investigations, the papillae also represent translocated parapodia and a similar naming (= buccal segment) should be used for the papillae-bearing region in Histriobdellidae. Notably, such a hypothesis would not only explain the formation of the head region in histriobdellids but would also cause a differing number of segmental ganglia. Thus, the ganglion previously considered to be the first one, would now be at the second position, right after the buccal segment (see also Fig. 6). The separation of the two ganglial halves could be seen as a result of reduced body size but relatively less reduced jaw-size in Histriobdellidae, forcing these ganglia apart and positioning them halfway along the long muscular pharynx instead of posterior to the latter. Notably, buccal structures can be found in different families among Eunicida and are often considered being an autapomorphic character for this taxon (Tilic et al, 2022; Kuhl et al., 2022). Nevertheless, detailed investigations also using developmental stages are lacking to confirm a homology of the histriobdellid buccal papillae and buccal structures in other Eunicida.

**Fig. 6:**
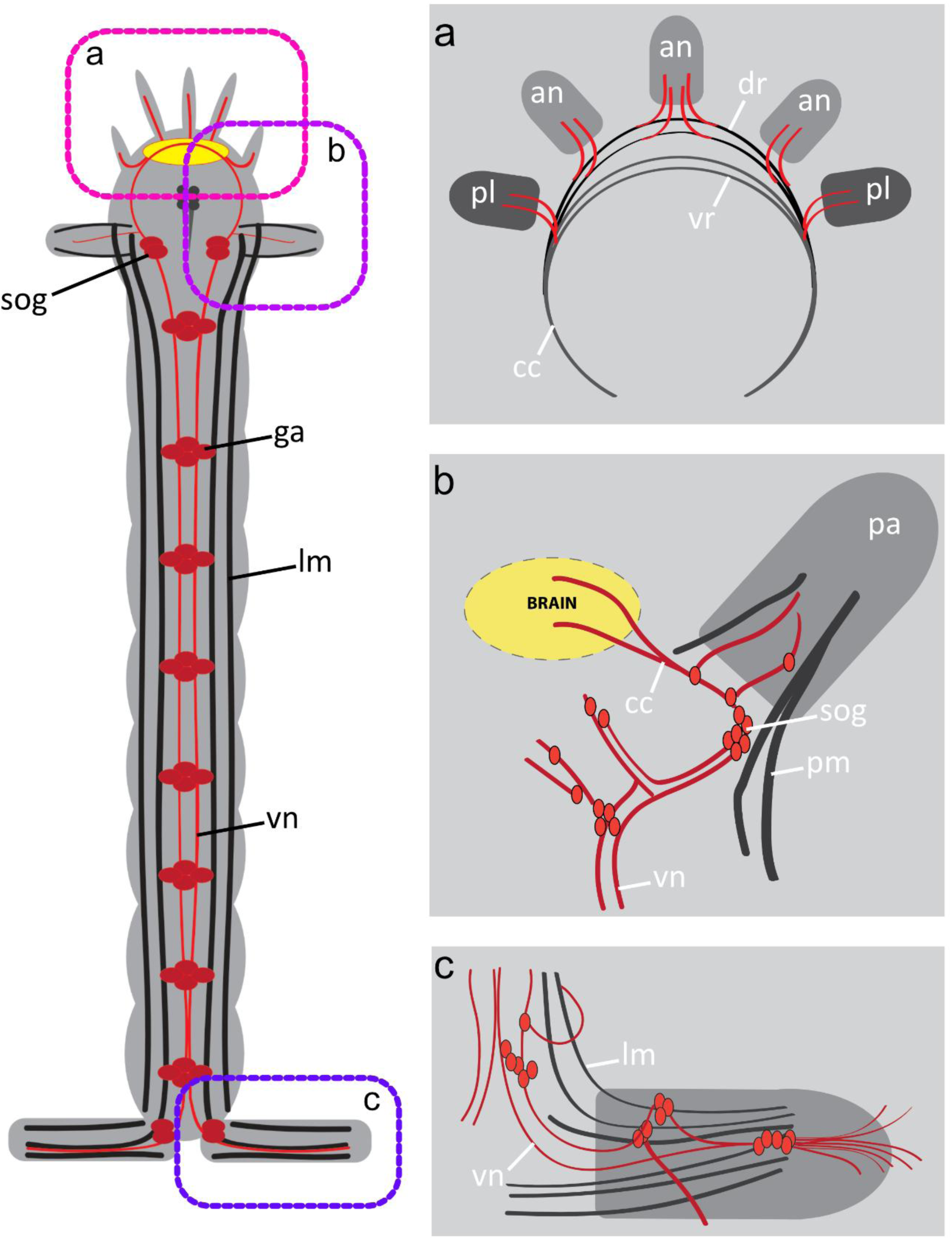
Schematic overview of the neuronal scaffolds in the anterior (a), buccal (b) and posterior (c) region of adult Histriobdellidae. Anterior is up. The red circles represent somata, red lines indicate main neurite bundles and black lines show the main pattern of muscle fibres. an, antennae; cc, circumesophageal connectives; dr, dorsal root of the cc; ga, ganglion; lm, longitudinal muscle bundles; pa; adhesive papilla; pm, muscle bundles within papillae; sog, subesophageal ganglion; vn, ventral nerve cord; vr, ventral root of the cc.

Another remarkable characteristic of the histriobdellid papillae is the presence of distinct glandular organs. This so-called duo-gland organ is known to bear two (adhesive and releasing) types of gland cells. As described by Gelder and Tyler (1986), the duo-gland organ is composed of cells producing acidophilic or “releasing” glycoproteins responsible for attachment and detachment of the papillae during locomotion. Comparable glands secreting various substances are also known for other annelid groups, such as interstitial Annelida (Worsaae et al., 2021) or Clitellata (e.g., Weigl, 1994). Furthermore, as shown by our investigations in histriobdellids, the longitudinal muscles of the appendage run along the gland cell necks, seem to terminate halfway along the latter necks and possibly make loop-shaped closures around the neck of the glands. Such an arrangement would allow controlling the release of secretions within the gland and is also known for the anterior end of Lobatocerebridae (Kerbl et al., 2015). Nevertheless, the exact arrangement of the gland neck muscles needs to be reinvestigated and is not described in detail by our data.

### The posterior appendages

Similar to the anterior adhesive appendages, the foot-like posterior ones bear prominent glandular distal openings. The respective glandular cells are known to possess a similar duo-gland system as shown for the anterior parapodia (Gelder & Seth Tyler, 1975). However, our muscular stainings show a different anatomy in the posterior appendages, with the glandular bundle proximally lacking supporting glandular sheet muscles as shown for the parapodial gland musculature in the anterior appendages. A similar strong tubulinergic signal of the gland necks can also be found in comparable gland systems in Lobatocerebridae (Kerbl et al., 2015) or Diurodrilidae (Worsaae & Rouse, 2008), and is caused by a more frequent occurrence of tubulins in the gland cell membranes. Additionally, the muscular origin differs since most muscular fibres supporting the posterior appendage extends from the trunk body wall musculature. Only very few muscle bundles such as the prominent bracing muscles are exclusive to the appendages. Same is true for the neuronal innervation. As already stated previously (see Gelder & Jennings, 1975) and verified by our investigations, the main neurite bundles innervating the posterior appendage come from the 8^th^ ganglion. Nevertheless, the posterior cirrus is solely innervated by neurites branching off one of the main neurite bundles and the posterior gland system gets innervation from neurites originating in an additional ganglion situated within the posterior appendage.

Accordingly, the muscular and neuronal innervation is highly different from the patterns observed for the anterior (parapodial) appendages. Hence, a prominent split of the ventral nerve cord running into the posterior appendages as well as disconnected ventral cord ganglia – as observed herein – has not been reported for other taxa so far. When closely observing innervation of the posterior appendages, it seems likely that – mainly based on the neuronal as well as muscular scaffold extending from the trunk structures – the proximal part of the posterior appendage including the cirrus might have evolved from parapodia-like structures. Such a hypothesis is furthermore supported by the presence of longer and numerous cirri at the proximity of the posterior appendage in other Histriobdellidae (Harrison, 1928). Additionally, the unique neuronal innervation of the distal part of the appendage with its origin in the 9^th^ ganglion as well as the presence of prominent gland structures, supports the distal appendage part as being evolved from pygidial lobes or cirri. According to the neuronal innervation of the glandular “feet”, a comparable scaffold can be observed in the interstitial genera *Diurodrilus* and *Saccocirrus*, supporting the latter hypothesis (Worsaae & Rouse, 2010; Worsaae et al., 2021). Furthermore, similar innervation patterns can be found for the pygidial cirri of e.g., Nereididae, Phyllodocidae as well as other errant Annelida (Starunov et al., 2015; Starunov, 2019). Those annelid posterior-most appendages are marked by a prominent muscular and neuronal innervation, which – best shown for the neuronal meshwork – represents an extension of the ventral nerve cord. According to these comparative findings and the anatomical details observed for *Histriobdella*, an evolutionary transformation of the last parapodial pair as well as the closely situated pygidial lobes into the unique posterior appendages of Histriobdellidae seems plausible. Nevertheless, further developmental studies are necessary to strengthen this hypothesis.

## Conclusion

Our analyses employ both molecular and morphological approaches for setting up hypotheses on the evolution of the enigmatic Histriobdellidae.

Accordingly, our molecular analyses support histriobdellids as being the sister group to the eunicid family Dorvilleidae. Although the entire body plan differs from that of Dorvilleidae or other members of the Eunicida, several comparable structures are observable that allow to set up homology assumptions. Hence, our data verify the homology of the three anterior-most head appendages to antennae, and support a homologization of the ventro-lateral appendages with the palps found in Dorvilleidae and other annelid families. Moreover, our investigations show that the adhesive anterior locomotory appendages (papillae) of *Histriobdella* should be referred to as being parapodia placed on a first buccal segment, previously not counted as such and differing from other Eunicida families all lacking parapodia on the first segment. Lastly, the adhesive posterior-most appendages show characters of both parapodial and pygidial structures, suggesting that the distalmost glandular part should be viewed as pygidial lobes whereas the proximal part may be seen as parapodial. In summary, our comprehensive analyses provide a phylogenetic placement for a so far unplaced group of Annelida and present important insights into the evolutionary plasticity of annelid body appendages and body plans.

## Declarations

## Ethics approval and consent to participate

No permits or authorizations were required to collect specimens used in the study.

## Consent to publish

Not applicable.

## Availability of data and materials

The datasets analysed during the current study are available from the corresponding authors on request. All data needed are included in the paper and the additional files.

## Authors’ contributions

NB and CH designed the project and drafted the manuscript. Data acquisition and analysis was done by all authors, and the final manuscript was read and revised by all authors.

## Authors’ information

Not applicable.

## Acknowledgements

Confocal imaging was performed at the Sars centre (Bergen, Norway). Scanning electron microscopy was conducted at the ELMILAB (University of Bergen) with help of Irene Heggstad. Molecular work was done at the DNA lab (University of Bergen) with help of Louise Lindblom. We thank Daniel Thiel (now University of Exeter) for aliquots of the anti-CFLRFamide antibody. Furthermore, we thank Irma Vila (University of Chile) for samples of *Stratiodrilus*.

## Open Access

This article is distributed under the terms of the Creative Commons Attribution 4.0 International License (http://creativecommons.org/licenses/by/4.0/), which permits unrestricted use, distribution, and reproduction in any medium, provided you give appropriate credit to the original author(s) and the source, provide a link to the Creative Commons license, and indicate if changes were made. The Creative Commons Public Domain Dedication waiver (http://creativecommons.org/publicdomain/zero/1.0/) applies to the data made available in this article, unless otherwise stated.

---

**Suppl. Fig. 1:** Best Maximum Likelihood tree resulting from a combined data set including COI, 16S rDNA, 18S rDNA and 28S rDNA. The tree topology include 53 taxa out of almost all families within the Eunicida (except Hartmaniellidae). The colour-coded drawings illustrate the shape of the jaw apparatuses in the respective families.

**Suppl. Fig. 2:** Table summarizing all taxa used for the phylogenetic analyses, with accession numbers and references provided.

