## Supplementary figures and images for "New insights into the evolution of Charlie Chaplin worms (Histriobdellidae, Eunicida, Annelida)"

### Supplemental Fig. 1

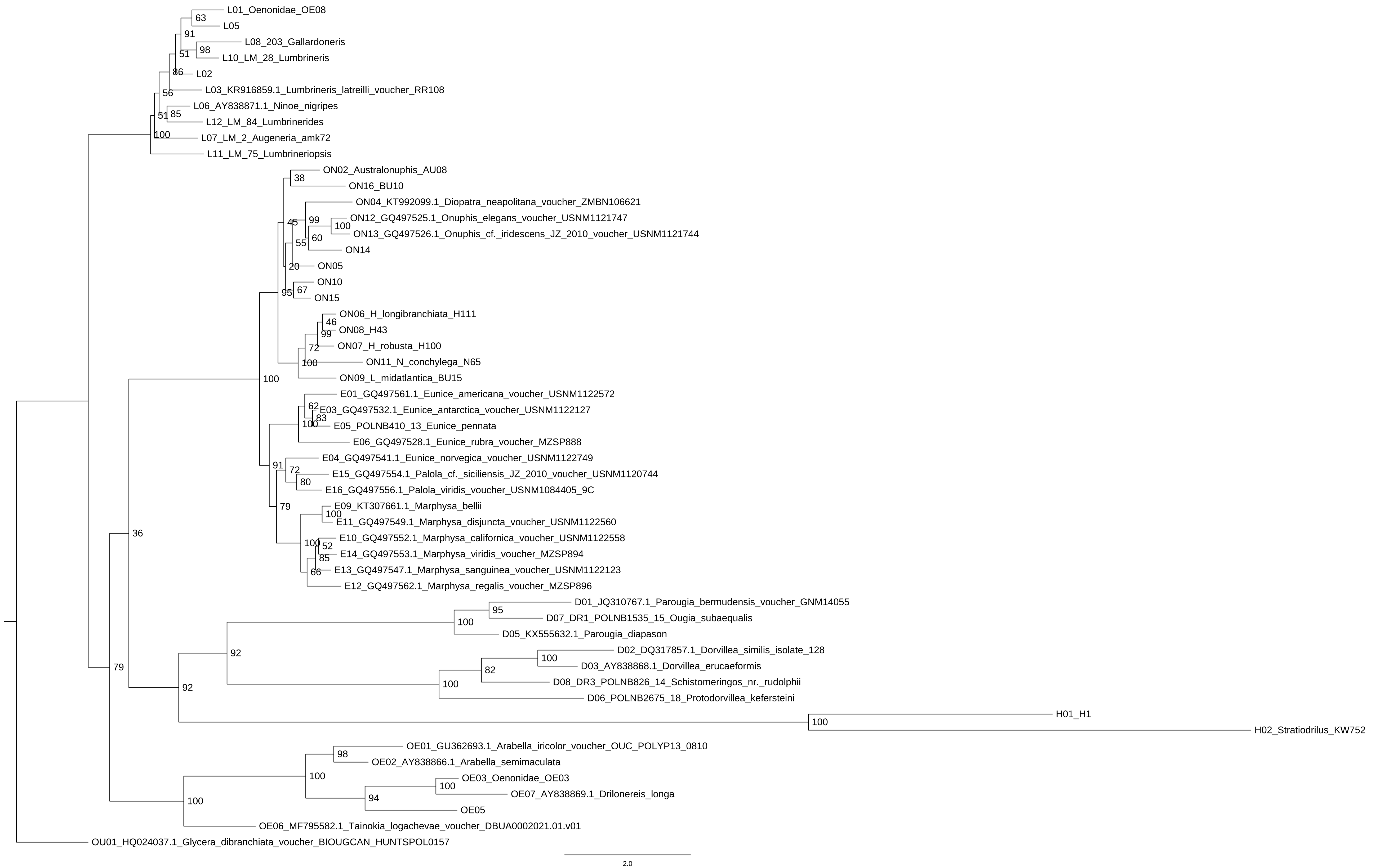
